# Bone morphogenetic protein signaling regulates Id1 mediated neural stem cell quiescence in the adult zebrafish brain via a phylogenetically conserved enhancer module

**DOI:** 10.1101/787804

**Authors:** Gaoqun Zhang, Marco Ferg, Luisa Lübke, Masanari Takamiya, Tanja Beil, Victor Gourain, Nicolas Diotel, Uwe Strähle, Sepand Rastegar

## Abstract

In the telencephalon of adult zebrafish, the *inhibitor of DNA binding 1* (*id1*) gene is expressed in radial glial cells (RGCs), behaving as neural stem cells (NSCs), during constitutive and regenerative neurogenesis. Id1 controls the balance between resting and proliferating states of RGCs by promoting quiescence. Here, we identified a phylogenetically conserved cis-regulatory module (CRM) mediating the specific expression of *id1* in RGCs. Systematic deletion mapping and mutation of conserved transcription factor binding sites in stable transgenic zebrafish lines reveal that this CRM operates via conserved *smad1/5 and 4* binding motifs (SBMs) under both homeostatic and regenerative conditions. Transcriptome analysis of injured and uninjured telencephala as well as pharmacological inhibition experiments identify a crucial role of bone morphogenetic protein (BMP) signaling for the function of the CRM. Our data highlight that BMP signals control *id1* expression and thus NSC proliferation during constitutive and induced neurogenesis.

## Introduction

In contrast to the mammalian adult brain, which contains only two main neurogenic regions that are both located in the forebrain and have rather limited ability to generate new neurons or to repair an injury, the brain of adult zebrafish contains numerous proliferative regions ^1-3^. These are distributed throughout different subdivisions of the brain and show high reactivation and repair capability upon lesion during adulthood ^4-7^.

The ventricular zone of the adult zebrafish telencephalon is the most extensively studied neurogenic region in this context. This region produces new neurons, which integrate into existing neural networks throughout the lifetime of the animal ^1, 2, 8, 9^. It is densely populated by the cell bodies of radial glia cells (RGCs), which are the neural stem cells (NSCs) of the adult telencephalon ^9, 10^. The cell bodies of the RGCs extend two processes: a short one to the ventricular surface and a long one that crosses the brain parenchyma to reach the pial surface ^9, 11^. Under homeostatic conditions, the majority of RGCs are quiescent (type 1) ^9, 12, 13^ and express typical neural stem cell markers such as glial acidic fibrillary protein (Gfap), brain lipid binding protein (Blbp) and the calcium-binding protein ß (S100ß) ^9, 10, 14^. Only a very low percentage of RGCs proliferate (type 2) and express the proliferative cell nuclear antigen (PCNA). This latter population can give rise to new neuroblasts (type 3 cells), either through asymmetric division or direct conversion ^15^.

A stab wound injury inflicted upon the telencephalon of adult zebrafish leads to an increase in NSC proliferation from 48 hours to 13 days post-injury ^16^ and a concomitant sustained production of new neurons that migrate from the ventricular layer to the injury site to replace the damaged/lost neurons ^17-20^. Remarkably, three weeks after the brain injury, the fish feed and breed normally, barely exhibiting histological traces of the traumatic damage ^17-21^. To quickly and efficiently replace dying/dead neurons, the number of RGCs entering the cell cycle and starting proliferation is greatly increased during regenerative neurogenesis ^15, 17, 19, 20^. Inflammatory signaling molecules cause expression of Gata3, a zinc finger transcription factor necessary for proliferation of RGCs, neurogenesis and migration of newborn neuroblasts ^5, 22^. In order to maintain a continuous supply of new neurons and to simultaneously prevent the exhaustion of the adult NSC pool, a tight control between quiescence, proliferation, differentiation and self-renewal of the RGCs is crucial. In a screen for transcriptional regulators expressed in the telencephalon of adult zebrafish, we previously identified the *helix-loop-helix factor id1* (*inhibitor of DNA binding 1*) as a key player of both homeostatic and regenerative neurogenesis ^23-25^. *Id1* is mostly expressed in quiescent RGCs, and its expression is up-regulated in the ventricular zone upon telencephalic injury. Forced expression of *id1* causes quiescence of neural stem cells, while *id1* knockdown increases the number of proliferating RGCs ^25^. *Id1* reduces cycling NSCs both during constitutive neurogenesis and after the initial wave of induction of proliferation in reactive neurogenesis ^25^. Our data argue for a role of *id1* in maintaining the balance between dividing and resting neural stem cells by promoting RGC quiescence. We speculated that this might prevent depletion of the NSC pool.

Neither Notch signaling nor inflammatory signals appear to be involved in regulating *id1* expression in the adult telencephalon ^25^. These data raised the central question of how and by which specific signals *id1* expression is restricted to adult neural stem cells and is up-regulated after brain injury. To address these questions, we decided to investigate the mechanisms of transcriptional regulation of this gene during constitutive and regenerative neurogenesis.

We identified a phylogenetically conserved *id1* cis-regulatory module (CRM) that drives GFP expression in RGCs of the adult brain of transgenic zebrafish. This RGC specific CRM harbors transcription factor (TF) binding sites conserved between human, mouse and zebrafish *id1* homologues. Deletion mapping, mutations of the binding sites, as well as pharmacological inhibition and transcriptome analysis suggest a role for the BMP pathway in controlling *id1* expression in RGCs both during constitutive and regenerative neurogenesis.

## Materials and methods

### Zebrafish strains and maintenance

All experiments were performed on 6-12 months old AB wild-type (wt) fish or on the transgenic reporter line described in this paper. Husbandry and experiments on animals were performed in accordance with the German animal protection standards and were approved by the Government of Baden-Württemberg, Regierungspräsidium Karlsruhe, Germany (Aktenzeichen 35-9185.81/G-272/12 and 35-9185.81/G-288/18 “Adulte Neurogenese”).

### Identification and cloning of putative cis-regulatory modules (CRMs)

Identification of CRMs driving *id1* expression in zebrafish was based on their conservation in comparison to other fish species. We utilized Ancora (http://ancora.genereg.net) to select sequences for functional analysis. Ancora represents a database of highly conserved noncoding elements (HCNEs) that are identified by scanning pairwise BLASTZ net whole-genome alignments with different similarity parameters ^26^. Criteria to be selected for functional testing were 80% sequence identity in a 50 bp window in all of the species *Oryzias latipes, Gasterosteus aculeatus* and *Tetraodon nigroviridis* within a 100 kb window around the *id1* locus. Genomic coordinates in putative CRMs chosen to test for regulatory potential were PCR-amplified from genomic DNA. Amplicons were subcloned into pCR™8/GW/TOPO® (Invitrogen) to create entry vectors for subsequent cloning into the Tol2-GFP-destination vector pT2KHGpzGATA2C1 as described by ^27^. These constructs were used to generate stable zebrafish transgenic lines. The sequences of all CRMs are provided in the supplementary data file S1.

### Mutation and deletion of different binding sites in the *id1* core sequence

Individual transcription factor binding sites were mutated by converting the core sequence of the binding site as defined by MatInspector to a stretch of thymidines. The approach was PCR-based by employing primers designed to include the desired change. Deletions were created using a similar methodology, a PCR-based approach using tailed primers designed to overlap with and anneal to the opposite strand of the adjoining region. The sequences of all primers used in this study can be found in supplementary data file S2.

### Injection of plasmids

For the generation of transgenic fish via a Tol2 based approach ^28^, 50 ng/µl plasmid DNA was injected. The injection solution was prepared by adding 1% of phenol red and 30 ng/µl *Tol2* mRNA. After injection the embryos were incubated at 28°C until they reached the desired stage. Embryos expressing GFP at 24 hpf were selected and the expression patterns documented. Identified F0 were out-crossed with wt fish to obtain stable progenies that express the transgene in the F1 generation. Each reporter construct was tested in at least 3 independent transgenic lines.

### Stab wound and chemical treatment of adult zebrafish

The stab wound injury was always inflicted in the left telencephalon hemisphere while the contralateral right hemisphere was kept intact and served as a control. 3 to 7 animals per transgenic line were tested for GFP induction upon telencephalon injury.

The stab wound procedure was performed as described ^21^. For the treatment, 9 month old *Tg(id1-CRM2:GFP)* fish were bathed in 300 ml fresh fish water containing 600 µl of a 10 mM DMH1 (Tocris, Wiesbaden-Nordenstadt, Germany) stock solution (final concentration 20 µM) for seven days. As a control, 600 µl of DMSO were added to 300 ml of fresh fish water. Every morning the fish were fed with regular adult fish food and the DMH1 or DMSO solution was changed every two days. Stab wounds were inflicted as described on the second day of treatment and the fish sacrificed for analysis 5 days after the injury. All experiments were carried out independently at least three times.

### Preparation of adult zebrafish brains, *in situ* hybridization, immunohistochemistry, imaging and quantification

Brain dissection, *in situ* hybridization and immunohistochemistry were performed as described in ^1, 21^. Primary antibodies used in this study include chicken anti-GFP (1:1000, Aves labs, Davis, CA), mouse anti-PCNA (1:500, Dako, Agilent, Santa Clara, CA) and rabbit anti-S100 (1:400, Dako). Secondary antibodies were conjugated with Alexa fluor dyes (Alexa series) and include anti-GFP Alexa 488, anti-mouse Alexa 546 and anti-rabbit Alexa 633 (1:1000, Invitrogen, Waltham, MA). Pictures of *in situ* hybridized sections were acquired with a Leica compound microscope (DM5000B). For imaging and quantification immunohistochemistry brain slices mounted in Aqua-Poly/Mount (Cat No. 18606-20, Polysciences, Inc) with #1.5 thickness coverslips were imaged with a laser scanning confocal microscope Leica TCS SP5. To obtain single-cell resolution images, an HCX PL APO CS x63/1.2NA objective was used with the pin-hole size set to 1-airy unit. Fluorescent images for GFP, PCNA and S100β were acquired sequentially in 16-bit color depth with excitation/emission wavelength combinations of 488 nm/492-550 nm, 561 nm/565-605 nm and 633 nm/650-740 nm, respectively. Pixel resolution for XY and Z planes are 0.24 and 0.50 µm, respectively. For individual brain samples, at least 3 transverse slices cut with a vibratome (VT1000S, Leica) at different anterior-posterior levels representing anterior, posterior an intermediate telencephalic regions were imaged.

In order to quantify the changes in *bambia, id3* and *smad5* mRNAs following brain injury at 5 days post lesion, three telencephalic pictures at the injury site were taken for each of the genes. Then, using image J, the pictures were processed setting up a threshold for the staining intensity and quantification of the up-regulated area was performed along the control and stab-wounded ventricular zone from the dorsomedial to the dorsolateral part of the telencephalon. The fold induction between the injured versus control ventricular zone was subsequently calculated.

## Image analysis

Confocal brain images were opened with Fiji/ImageJ software ^29^ as composite hyperstacks to manually evaluate colocalization of GFP, PCNA and S100β proteins. Colocalization of fluorescent signals was assessed by at least two experimenters. For quantifications, 3 sections per brain were analyzed. Cells were counted in dorsomedial and the dorsolateral ventricular zone.

## Statistical Analysis

For quantifications of *id1-CRM2:GFP* and derivative constructs, the number of cells was determined by counting the cells in Z□stacks of 50 μm thickness in 1 μm steps (×40 objective lens). Statistical significance was assessed by using R. Cells were always counted at the dorsomedial and dorsolateral regions of adult zebrafish telencephalon ventricular zone.

## Quantitative Realtime PCR

Total RNA was isolated from adult telencephalon using Trizol (Life Technology). First-strand cDNA was synthesized from 1 µg of total RNA with the Maxima First strand cDNA synthesis kit (Thermo Scientific) according to the manufacturer’s protocol. A StepOnePlus Real-Time qRT-PCR system (Applied Biosystems) and SYBR Green I fluorescent dye (Promega) were used. Expression levels were normalized using β-actin (Fig.7). The relative levels of mRNA were calculated using the 2^-ΔΔCT^ method. The primer sequences are listed in supplementary material Table S2.

### RNA sequencing and library preparation

For RNA-sequencing, total RNA was isolated from 3 injured hemispheres (5 days post injury) and 3 uninjured contralateral telencephalic hemispheres each to create injured and uninjured RNAseq libraries, respectively. Our data were generated from 3 biological repeats (as described ^25^). 2 mg of total RNA was used to prepare each of the 6 mRNAseq libraries. RNA sequencing data were retrieved from a previous data set ^25^. The data analysis was carried out as described ^30^ using the latest version of the zebrafish reference genome, assembly GRCz11. (https://www.ncbi.nlm.nih.gov/assembly/GCA_000002035.4/).

## Results

### Identification of a CRM mediating expression of *id1* in adult neural stem cells

To elucidate the mechanisms underlying *id1* expression in adult neural stem cells and its injury-induced up-regulation, we performed a systematic search for CRMs controlling expression of *id1* in the ventricular zone of the adult telencephalon. Via phylogenetic sequence comparison of the *id1* locus, we identified five conserved putative cis-regulatory modules (CRM1-5) upstream and downstream of the *id1* coding sequence (Fig. 1A). The identified conserved non-coding sequences were inserted in front of a *gata2* minimal promoter ^31^ coupled to a GFP reporter cassette ^27^ and introduced into the germ line of zebrafish to generate stable transgenic lines, *Tg(id1-CRMX:GFP)*, where X represents one of the five CRMs (Fig. 1B-F, H-L and H’-L’). All five CRMs mediated expression in 24-hour post fertilization (hpf) zebrafish embryos in somewhat overlapping but distinct and specific patterns (Fig. 1B-F). In the adult telencephalon (Fig. 1H-L and H’-L’), only *id1-CRM2* mediated specific GFP expression in the ventricular zone (white arrows, Fig. 1K and supplementary movie 1) in a pattern identical to the previously described GFP-tagged *id1* BAC (bacterial artificial chromosome) transgenic line, *TgBAC(id1:GFP)*)^25^ (Fig. 1M and M’), while CRM5 (Fig. 1H), CRM4 (Fig. 1I), CRM3 (Fig. 1J) and CRM1 (Fig. 1L) drove ectopic GFP expression presumably at the tela choridea and blood vessels (white rectangle, Fig. 1H, magnification in 1H’), in neurons (white rectangle, Fig. 1I, 1L and respective magnification in Fig. 1I’ and 1L’) and in blood vessels and presumptive oligodendrocytes/neurons (white rectangle, Fig. 1J and magnification in 1J’). The ventricular GFP positive cells resembled RGCs in morphology and co-expressed the RGC marker S100ß (96.9% ±2.3%), (Fig. 1K, K’, N, Q and R) which were shown to be the neural stem cells (NSCs) in the adult zebrafish telencephalon^9, 10^. GFP was predominantly expressed in S100ß^+^/PCNA^-^ type 1 cells (82.5 ±8.4%, n=3 telencephala; Fig. 1N-Q and R-U), while only a small number of GFP^+^/S100ß^+^ cells represented type 2 cells co-expressing PCNA (17.5 ±8.4%, n=3), a marker for cell proliferation GFP expression was also excluded from the rostral migratory stream (RMS), a highly proliferative domain in the telencephalon composed of type 3 progenitors (Fig. 1K, white arrowhead). These data show that GFP^+^ cells correspond in majority to quiescent type 1 RGCs. Furthermore, both GFP expression and intensity were increased in the left telencephalic hemisphere upon stab injury compared to right non-injured control hemispheres (Fig. 1R, U, V-W and Y-Z). After stab wounding the proportion of GFP^+^/S100ß^+^/PCNA^-^ type 1 and GFP^+^/S100ß^+^/PCNA^+^ type 2 stem cells was not altered in the injured left hemisphere relative to the right uninjured side, which is similar to the endogenous *id1* gene and the *id1* BAC transgenic line^25^. Additionally, GFP was predominately found in quiescent cells (Fig. 1W-X). Thus, the *id1-CRM2* drives expression of GFP in the RGCs in a pattern identical to the endogenous *id1* gene, and this expression is inducible by injury.

**Figure 1:**
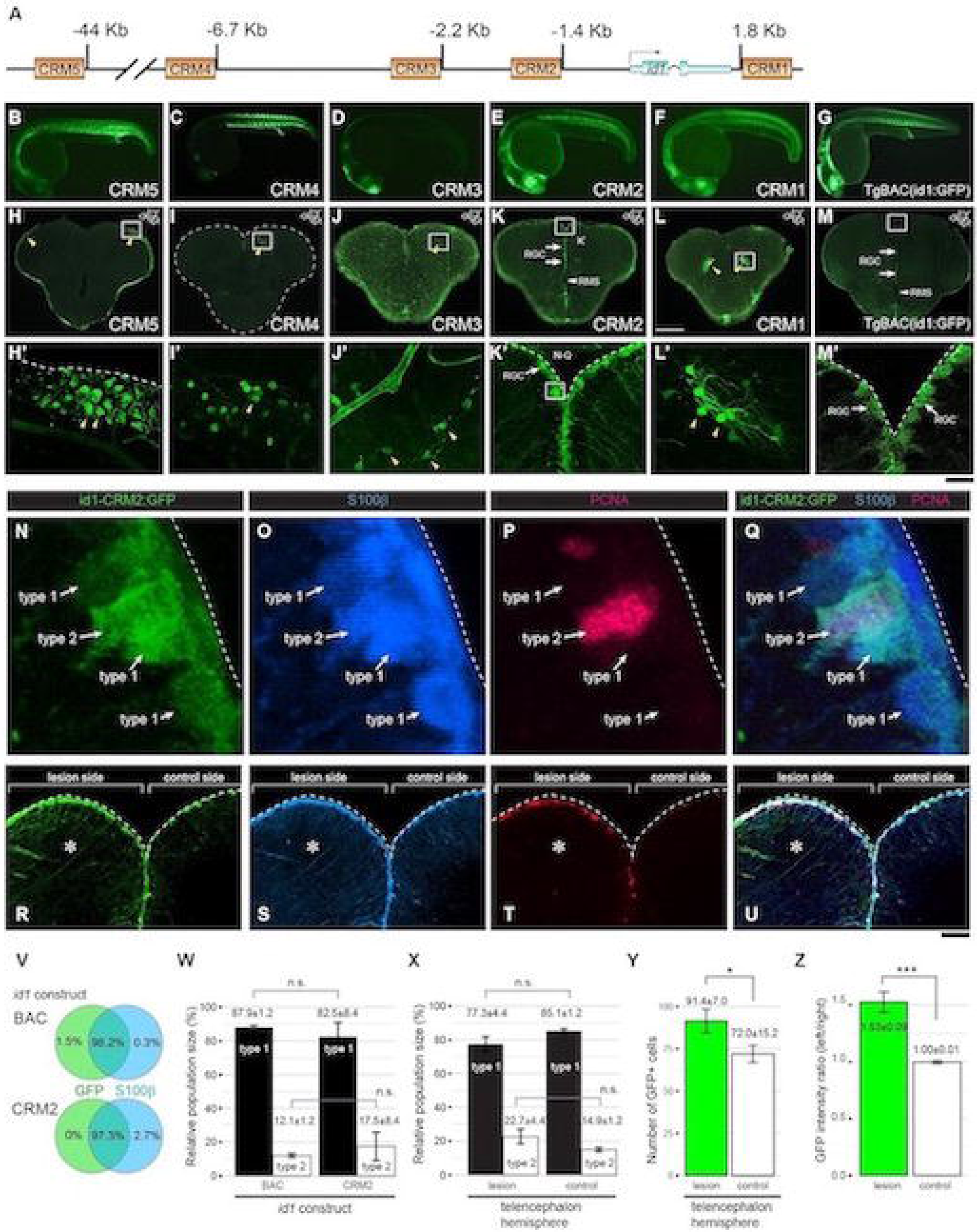
The *id1* Cis-regulatory module 2 (CRM2) drives ventricular expression and responds with increased expression to stab injury of the adult zebrafish telencephalon. (A) Homology-based search for cis-regulatory modules (CRMs). Schematic representation of the *id1* locus with putative CRMs 1-5 highlighted by yellow rectangles. (B-M’) Stable GFP reporter expression **of** *id1*-CRM5 (B and H-H’), *id1*-CRM4 (C and I-I’), *id1*-CRM3 (D and J-J’), *id1*-CRM2 (E and K-K’), *id1*-CRM1 (F and L-L’) are shown in comparison with the control *TgBAC(id1:GFP)* (G and M-M’) for 24 hpf embryos (B-G) and adult telencephala (H-M’). (H-M) GFP reporter expression analyzed in transverse sections from the middle part of the adult telencephalon (location of section schematically indicated in the upper right corner). The *Tg(id1-CRM2:GFP)* transgenic line (K) recapitulates GFP expression of the *TgBAC(id1:GFP)* line (M) with strong expression in the ventricular zone (white arrows) and absence of expression in the rostral migratory stream (RMS, white arrowheads). The yellow arrowheads indicate ectopic GFP expression in the tela choroidea (H-H’), cells with appearances of neurons (I-I’ and L-L’) and oligodendrocytes (J-J’). Rectangles in the panels H-M represent the region magnified in H’-M’, respectively. Dashed lines indicate the boundary of the telencephalon. (N-Q) Magnified views of RGCs highlighting type 1 and type 2 RGCs identified by expression of *id1-CRM2:GFP* (N), S100ß (O) and PCNA (P; all merged in Q). (R-U) Transverse view of a 5 days post injury telencephalon showing the expression of *id1-CRM2:GFP* (R), S100ß (S) and PCNA (T; all merged in U). The lesion site (left hemisphere) is marked by an asterisk. (V-W) Summary of colocalization analysis of GFP and S100ß expression for BAC and *CRM2- id1* constructs. (W) Relative population sizes of type 1 and type 2 RGCs for BAC and *id1-CRM2* constructs. (X-Y) Relative population sizes of type 1 and type 2 RGCs (X) and the number of GFP^+^ cells (Y) for *id1-CRM2* constructs, comparing lesioned and unlesioned control hemispheres. (Z) *id1-CRM2:GFP* intensity ratio between left and right hemispheres comparing undamaged telencephalon (control) and damaged telencephalon (lesion). Significance is indicated by asterisks: *, .01≤ *p* <.05; ***, *p* <.001. *n.s*.=not significant. Scale bars: 200 μm (H-M), 20 μm (B-G; H’-M’), 2 µm (N-Q) and 25 µm (R-U).

### Fine-mapping of the regulatory sequences mediating expression in RGCs

In order to map the core regulatory region of the *id1-CRM2* responsible for its specific activity in the RGCs of the telencephalic ventricular zone, a series of 5’and 3’ overlapping deletion variants of the reporter construct were generated and analyzed in stable transgenic lines (Fig. 2A and data not shown). The transcriptional activities of these mutant versions were first investigated by monitoring GFP expression in 24 hpf zebrafish embryos. We observed at least 3 independent transgenic lines per construct and only constructs driving strong GFP expression resembling *id1* embryonic expression were selected for further analysis in the adult brain (Fig. 2A and data not shown).

**Figure 2:**
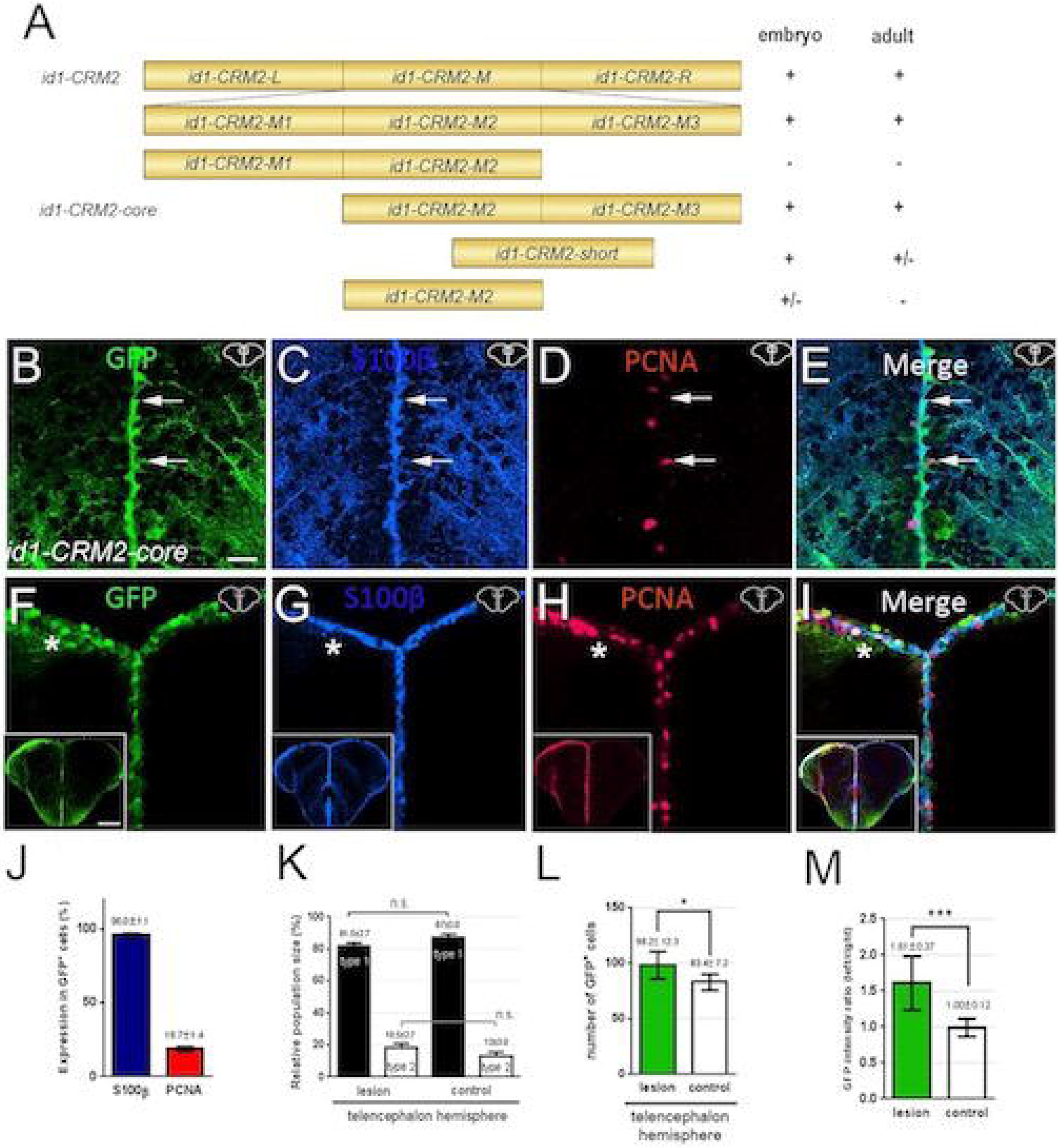
Deletion mapping of *id1-CRM2* identified a 157 bp core region, which confers RGC-specific expression in the adult telencephalon. (A) 5’and 3’ deletions of *id1-CRM2* analyzed for expression in zebrafish embryos and adult brains. Results are summarized on the right; + indicates specific GFP expression; +/- and – represent weak and absence of expression, respectively. (B-I) Immunohistochemistry of telencephalic transverse sections with antibodies against GFP (B, F), S100ß (C, G) and PCNA (D, H) (merged panels: E, I). Section levels and areas of magnification are indicated in the upper right-hand corner of the image. (B-E) White arrows show two RGCs. (E) The upper cell is GFP^+^/S100β^+^/PCNA^-^ (type 1 RGC) while the lower cell is GFP^+^/S100^+^/PCNA^+^ (type 2 RGC). (F-I) Upon stab wound injury the reporter construct expression is up-regulated. The left injured side is labelled with a white asterisk. (B-I) Section levels and areas of magnification are indicated in the upper right-hand corner of the image. (J) Quantification of PCNA and S100ß expression in *id1-CRM2:GFP* positive cells. (K) Relative population size of type 1 and type 2 RGCs in the control and lesioned hemisphere. (L, M) Quantification of GFP-positive cells and GFP intensity upon injury. Graphs showing the number of GFP-expressing cells (L) and the intensity ratio between left uninjured (control) and right injured hemispheres respectively (M). Bars: mean ± SD. Significance is indicated by asterisks: *, .01≤ *p* <.05; ***, *p* <.001. *n.s*.=not significant. Boxed in image in lower left-hand corner of (F-I) represents entire brain sections. *n*=3 animals (B-L), *n*=15 sections (M). Scale bar = 20 μm (B-I); 200 μm for Boxed image in lower left-hand corner of (F-I).

The deletion construct designed as *id1-CRM2-core* which contains a 157 bp long stretch of the CRM*2* sequence (Fig. 2A, chr11:18,706,838-18,706,994) drove expression in the ventricular zone of the adult telencephalon (Fig. 2B). Double labeling experiments of the transgenic line *Tg(id1-CRM2-core:GFP*) revealed that this shorter version of the CRM2 drives mainly expression in PCNA^-,^ S100β^+^ RGCs (Fig. 2B-E and J), as observed for the endogenous *id1* ^25^, *id1-BAC:GFP and id1-CRM2* (Fig. 1R, U and V). Moreover, this short sequence responded to injury by increased expression and intensity of GFP (Fig. 2F-I and L, M; n=3 telencephala). Thus, our deletion approach led to the identification of a 157 bp sequence that appears to harbor all relevant sequences to drive expression in neural stem cells and to respond to injury. Remarkably, this sequence also proved sufficient to drive GFP expression in the brain, eye, somites, midline and urogenital opening in 24 hpf embryos (supplementary Fig. 1) indicating that both embryonic and adult regulatory signals act through this CRM.

### *id1-CRM2-core* is structurally and functionally conserved between fish and human

Sequence analysis of zebrafish *id1-CRM2-core* with the *MatInspector* software ^32^ showed that this regulatory module harbors transcription factor (TF) binding sites for Forkhead box protein A2 (FoxA2), Cyclic AMP response element binding protein (CREB), Homeobox domain transcription factor (Pknox), and Early growth response gene 1 (Egr1), as well as two *smad binding motifs (SBMs)* (Fig. 3A). The entire core sequence displays a high degree of conservation between zebrafish, mouse and human homologues. We thus tested whether the sequence is also functionally conserved by constructing a transgene harboring the human version of the zebrafish CRM2-core referred to as *Hsid1-CRM2-core*. After stably introducing this transgene into the germ line of zebrafish, we analyzed expression in the adult telencephalon (Fig. 3B-E). The human sequence also mediated expression in the ventricular zone, in type 1 progenitors corresponding to quiescent RGCs (PCNA^-^, S100ß^+^), similar to the zebrafish sequence (Fig. 3J and Fig. 2J, respectively). Moreover, when a brain injury was inflicted in one hemisphere of the telencephalon the *Hsid1-CRM2-core* carrying transgene also responded to the stab wound by increased expression at 5 days post-lesion (dpl) (Fig. 3F-I and L, M; n=3 telencephala). Taken together, these results show that the mechanism of *id1* regulation appears to be conserved from fish to human, suggesting that the mechanisms underlying the control of neurogenesis are very similar despite the remarkably different abilities to repair lesions in the adult brain among these phylogenetically distant vertebrate species.

**Figure 3:**
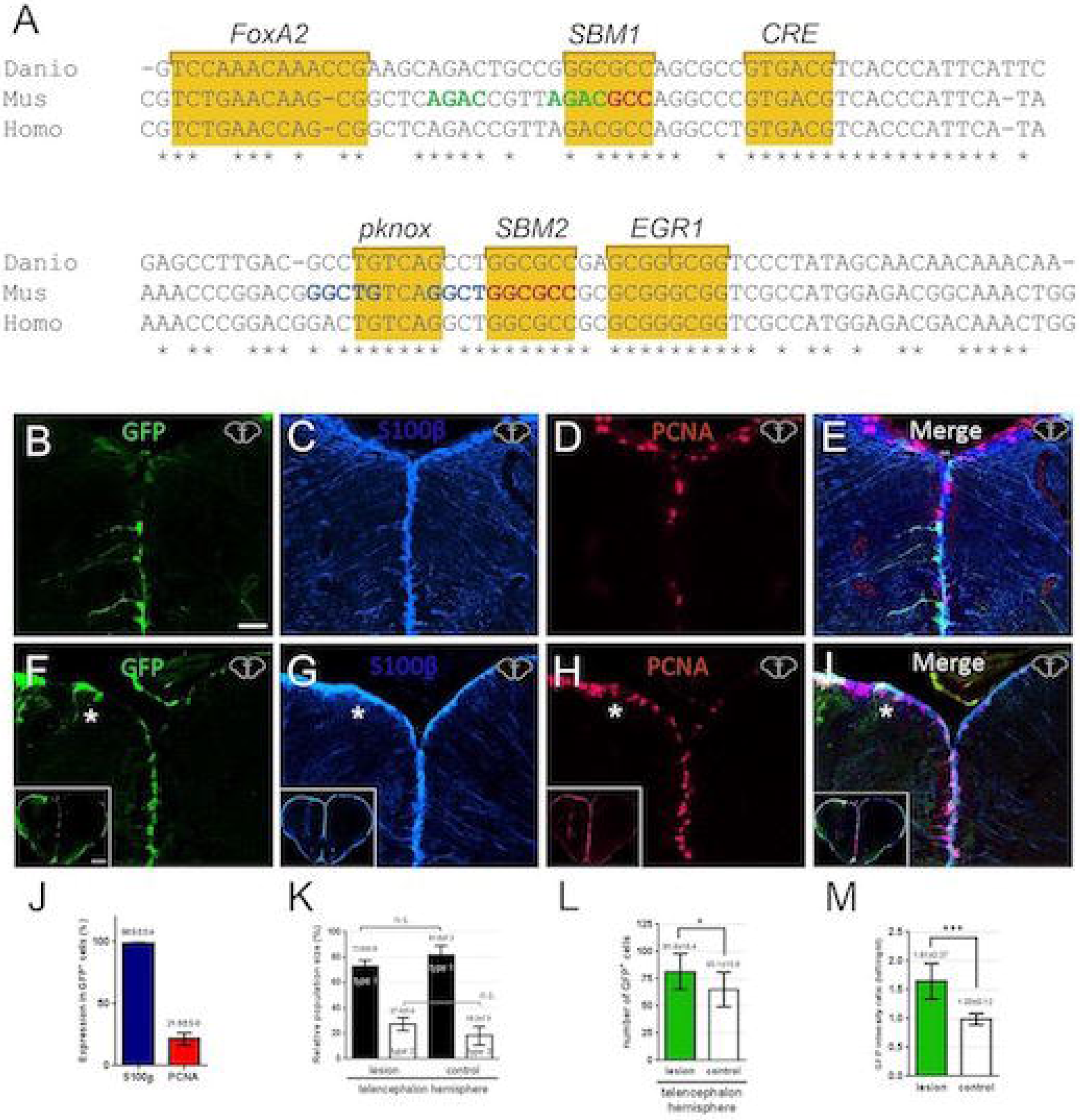
Conservation of the zebrafish *id1-CRM2* core sequences and its function across evolution. (A) Sequence comparison of zebrafish *id1-CRM2-core* (Danio) with human (Homo) and mouse (Mus) orthologous sequences. Conserved nucleotides are indicated with an asterisk. Conserved motifs are outlined by yellow boxes comprising putative DNA recognition sequences for the transcription factors FoxA2, Smad (SBM1 and 2), CRE binding protein (CREB), Pknox and EGR1. Nucleotide sequences in green, red or blue correspond to previously identified sequences in the mouse *id1* orthologue: *Smad binding element (SBE)*, a Smad 1/5 binding site and a binding site for an unknown binding protein, respectively. (B-I) Immunohistochemistry of telencephalic transverse sections with antibodies against GFP (B, F), S100ß (C, G) and PCNA (D, H) (merged panels: E, I). (B) Expression of GFP in RGCs at the telencephalic ventricular zone driven by the human *id1* regulatory sequences in the zebrafish adult telencephalon. (F) Expression of the human *Tg(Hsid1-CRM2)* driven reporter construct is up-regulated upon stab wound injury. The injured telencephalic hemisphere is labeled with a white asterisk. (J) Quantification of PCNA and S100ß expression in GFP^+^ cells in the *Tg(Hsid1-CRM2)* line. (K) Relative population size of type 1 and type 2 RGCs in the control and lesioned hemispheres. The proportion of GFP^+^/S100β^+^/PCNA^-^ type 1 and GFP^+^/S100^+^/PCNA^+^ type 2 stem cells is not altered in the injured hemisphere relative to the control hemisphere of the telencephalon. (L, M) Quantification of GFP^+^ cells upon injury. The number of GFP-expressing cells (L) and the intensity ratio between left and right hemispheres comparing undamaged hemisphere (control) and damaged telencephalic hemisphere (M) are both increased following injury. Bars: mean ± SD. Significance is indicated by asterisks: *, .01≤ *p* <.05; ***, *p* <.001. *n.s*.=not significant. Boxed in image in lower left-hand corner of (F-I) represents entire brain sections. *n*=3 animals (B-L), *n*=15 sections (M). Scale bar = 20 μm (B-I); 200 μm for Boxed image in lower left-hand corner of (F-I).

### BMP response elements are required for *id1-CRM2* activity in the zebrafish adult telencephalon

The central region (chr11:18,706,875-18,706,953) of the *id1-CRM2-core*, situated between the *foxA2* and *egr1* binding sites and containing two *SBMs* (Fig. 3A and 4A), is similar to a previously identified *BMP response element* (*BRE*) of the mouse and human *id1* gene ^33, 34 35^. Because *id1* is a direct target of the BMP signaling pathway ^36^ and smads transduce the BMP signal from the cytoplasm to the nucleus ^37^ (Fig. 6A), we tested whether this *BRE* is necessary for the function of *id1-CRM2* in the zebrafish adult telencephalon. To this aim we generated two *id1-CRM2* mutant variants, in which either the conserved 74 bp sequence covering the *BRE* was deleted (*id1-CRM2Δ74*) (Fig. 4B) or both *SBM1* and *SBM2* sequences were mutated (*id1-CRM2-mut-SBMs*) (Fig. 4C). GFP expression in the ventricular zone and in RGCs was abolished in transgenic lines carrying the mutant constructs (Fig. 4D-H (*id1-CRM2Δ74;* no GFP, expression in S100ß^+^ cells n=5) and (Fig. 4I-M (*id1-CRM2-mut-SBMs*, no GFP expression in S100ß^+^ n=6)) and no induction of GFP expression upon injury could be detected for both constructs (Supplementary Fig. 3A-D and E-H). These results show that the *BRE* located in the *id1-CRM2* is critical for the expression of *id1* in RGCs. Moreover, they suggest that BMP signaling may play a role in controlling *id1* expression in the telencephalon of the adult zebrafish.

**Figure 4:**
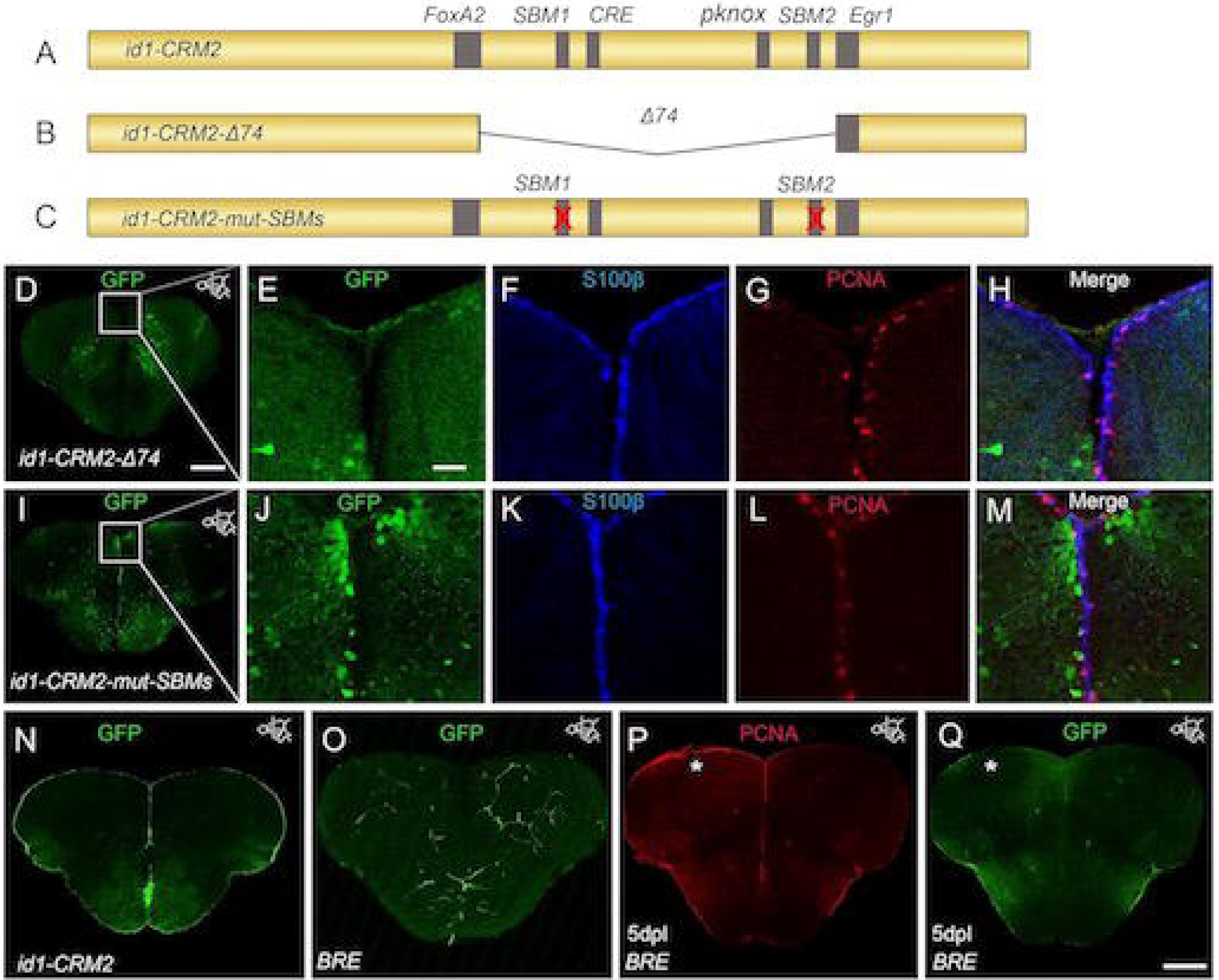
The conserved BMP response element in the *id1-CRM2* is crucial for correct expression of GFP reporter in the ventricular zone and RGCs. (A-C) Scheme showing mutated *id1-CRM2* reporter constructs: (A) *id1-CRM2* wt construct with putative TF binding sites indicated in grey, (B) *id1-CRM2-Δ74* construct which contains a deletion of a 74 bp stretch of the most conserved sequence in *id1-CRM2*, (C) *id1-CRM2-mut-SBMs* construct with mutation in the 2 SBMs (1 and 2) of *id1-CRM2.* (D) Deletion of the 74 bp stretch in the *id1-CRM2* abolished GFP expression in the ventricular zone. (E-H) Enlarged micrographs of (D). (E-H) Immunohistochemistry with GFP (E), S100ß (F) and PCNA (G) antibodies on telencephalic cross-sections of the *id1-CRM2-Δ74* transgenic line shows no GFP reporter expression in S100ß+ RGCs (H, merged view). (I-M) Mutations in *Smad binding motifs* (*SBM1* and *2*) abolished GFP expression in the ventricular zone. (J-M) Magnification of white-boxed region in (I). (J-M) Immunohistochemistry with GFP (J), S100ß (K) and PCNA (L) antibodies on telencephalic cross-sections of the *id1-CRM2-mut-SBMs* transgenic line show no colocalization between the RGC marker, S100ß, and GFP (M, merged view). (N) GFP expression driven by *id1-CRM2:GFP* reporter construct in the ventricular zone (control). (O-P) Immunohistochemistry of telencephalic cross-sections with GFP (O, Q) and PCNA (P). (O, Q) The *BRE* does not drive GFP expression in the RGCs (O) and is not inducible by telencephalic injury (Q). The left injured side is labelled with a white asterisk. Anteroposterior positions of transverse sections are indicated in the upper right-hand corner of each image. Scale bar = 20 μm (E, F, G, H, J, K, L, M); 200 μm (D, I, N, O, P, Q).

We next tested whether the well-characterized mouse *id1 BRE* ^33, 34^ that contains multiple *SBMs* would control expression in the telencephalon. A construct containing 2 tandem copies of the mouse *BRE* was shown to mediate GFP expression in all BMP signaling target tissues of zebrafish embryos ^38, 39^. Surprisingly, however, it did not show any activity in the RGCs of the ventricular zone of the adult telencephalon (Fig. 4O), and was unaffected by brain injury (Figure. 4P,Q). In the transgenic line *Tg(BRE:GFP)*, GFP expression is restricted to blood vessels of the adult brain (Fig. 4O, n=7), in contrast to the expressions driven by *id1-*CRM2 (Fig. 4N) and *id1-CRM2-core* (Fig. 2B), which are RGC-specific (compare Fig. 4N to Fig. 4O). This blood vessel expression was observed with two independent lines of the *BRE:GFP* described in ^38^ and ^40^. These findings suggest that in the adult zebrafish brain, the BMP pathway alone is critical but not sufficient to drive *id1* expression in the ventricular RGCs.

### *BRE* and additional transcription factors are necessary for full activity of *id1-CRM2* in neural stem cells

The observation that the two tandem copies of the mouse *BRE* ^33^ are not sufficient to drive reporter gene expression in RGCs (Fig. 4O) suggests that further sequences are required in addition to *SBMs* for activity of *id1-CRM2-core* in RGCs. Indeed, the *id1-CRM2-core* contains several other well-conserved binding sites for transcription factors surrounding the *SBMs* (Fig. 3). To test whether these neighboring sequences are necessary in addition to the *BRE*, each of these sites was individually mutated, and stable transgenic lines were generated with the resulting mutant constructs (Fig. 5A-E). While mutation in the *foxA2* and *egr1* binding sites had no effect on the expression pattern of GFP in the RGCs of the adult brain (Fig. 5B, E and 5P, Q), we observed a reduction in GFP expression for constructs with mutated *pknox* or *cre* binding sites (Fig. 5C, D and 5P, Q). Notably, we still observed a strong and specific response to stab injury of the telencephalon in these two mutant lines (Fig. 5F-J and 5P, Q and Fig. 5K-O and 5P, Q). This finding indicates that these sites together with *SBMs* are necessary for RGC-specific expression. However, since mutations of the individual *cre* (Fig. 5F-J and 5P, Q) and *pknox* (Fig. 5K-O and 5P, Q) sites did not affect the capacity to respond to injury, these sites appear not to be required for injury-induced expression via the BMP pathway.

**Figure 5:**
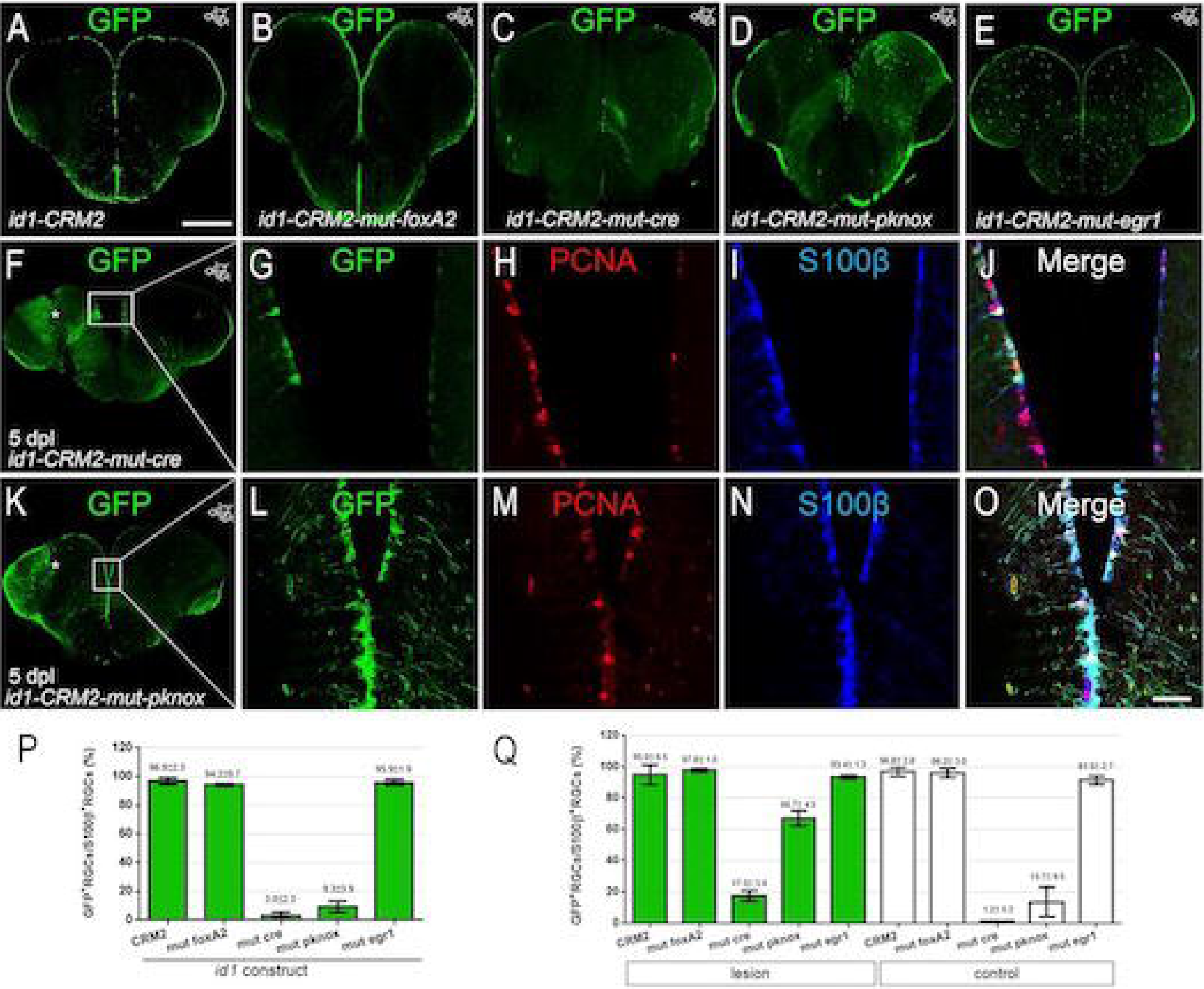
Cre and pknox binding sites in close vicinity to *SBM1* and *-2* are necessary for the full activity of *id1-CRM2*. (A-E) Immunohistochemistry with a GFP antibody on telencephalic cross-sections of the *id1-CRM2* transgenic line containing a mutation in the *foxA2* (B), *cre* (C), *pknox* (D) and *egr1* (E) binding sites, respectively. Mutation in *cre* (C) and *pknox* (D) binding sites leads to a reduction of GFP^+^ cells in the ventricular zone. (F, G, J) Expression of the *id1-CRM2-mut-cre* construct is inducible by brain injury (white asterisk, left injured hemisphere). (G-J) Magnification boxed area in (F). (K, L, O) Expression of the *id1-CRM2-mut-pknox* construct is induced upon brain injury (white asterisk, left injured hemisphere). (L-O) Magnification of boxed area in (K). Anterioposterior positions of transverse sections are indicated in the upper right-hand corner of each image. (P, Q) Percentage of GFP^+^/S100β^+^ RGCs over the total number of S100^+^ RGCs for *id1-CRM2* mutant constructs under homeostatic condition (P) and at 5 days post injury, comparing the injured hemisphere (green columns) and the contralateral uninjured hemisphere (white columns; Q) at the dorsomedial to the dorsolateral region of the telencephalon. Scale bar = 200 μm (A-F and K); 25 μm (G-J and L-O).

### *Id1-CRM2* expression is induced by the BMP pathway in response to injury

A crucial question is whether the increase in *id1-CRM2* mediated transcription at the ventricular zone upon injury involves BMP signaling, as suggested by the requirement of the BRE for CRM2 activity. To address this question, in a first step we analyzed deep sequencing data set of the transcriptomes generated from 5 days post lesion (dpl) hemispheres versus contralateral non-injured adult zebrafish telencephala ^25^. We discovered an increase in the transcription levels of several key genes involved in canonical BMP signaling (Fig. 6A, B) when compared to the control, uninjured side. Among those, we identified the *BMP receptor1aa (bmp1aa)*, the BMP receptor specific signal transducers *smad1 and smad5* as well as several direct target genes of BMP signaling, including *id1, id3, izts2a* and *bambia* (Fig. 6B). For genes which are significantly up-regulated (Fig. 6B) and which display a restricted pattern of expression in the telencephalon under homeostatic conditions (*bambia, smad5* and *id3*), the results were further verified by *in situ* hybridization (ISH) on telencephalic cross sections at 5 dpl (Fig. 6C-E). A distinct up-regulation of *bambia, smad5* and *id3* at 5 dpl in the injured left telencephalic hemisphere was detected on the sections when compared to the right uninjured hemisphere, where expression did not change (Fig. 6 C-E; n=4 telencephala per gene). The RNA sequencing and ISH data were confirmed by quantifying the staining intensity of *bambia, smad5* and *id3* following brain injury at 5 days post lesion at the injury site in comparison to the same region in the intact contralateral control hemisphere (Fig. 6 C’-E’).

To functionally manipulate BMP signaling during the response to injury, telencephala of adult fish were injured on the second day of treatment with 20 µM of the BMP signaling pathway inhibitor DMH1 (dorsomorphin homologue 1; ^41, 42^). The fish were analyzed at 5 dpl. As visualized by *in situ* hybridization (Fig. 7A, B, D, E) and by qPCR analysis (Fig. 7C, F), suppression of BMP signaling by exposure to DMH1 led to a significant loss of induction of the endogenous *id1* gene (Fig. 7B) and *id1-CRM2:GFP* reporter gene (Fig. 7E) upon injury of the telencephalon. We also observed a strong reduction in the basal expression of the *gfp* transgene and *id1* endogenous gene in the uninjured telencephalic hemispheres upon DMH1 treatment (Fig. 7C, F).

**Figure 6:**
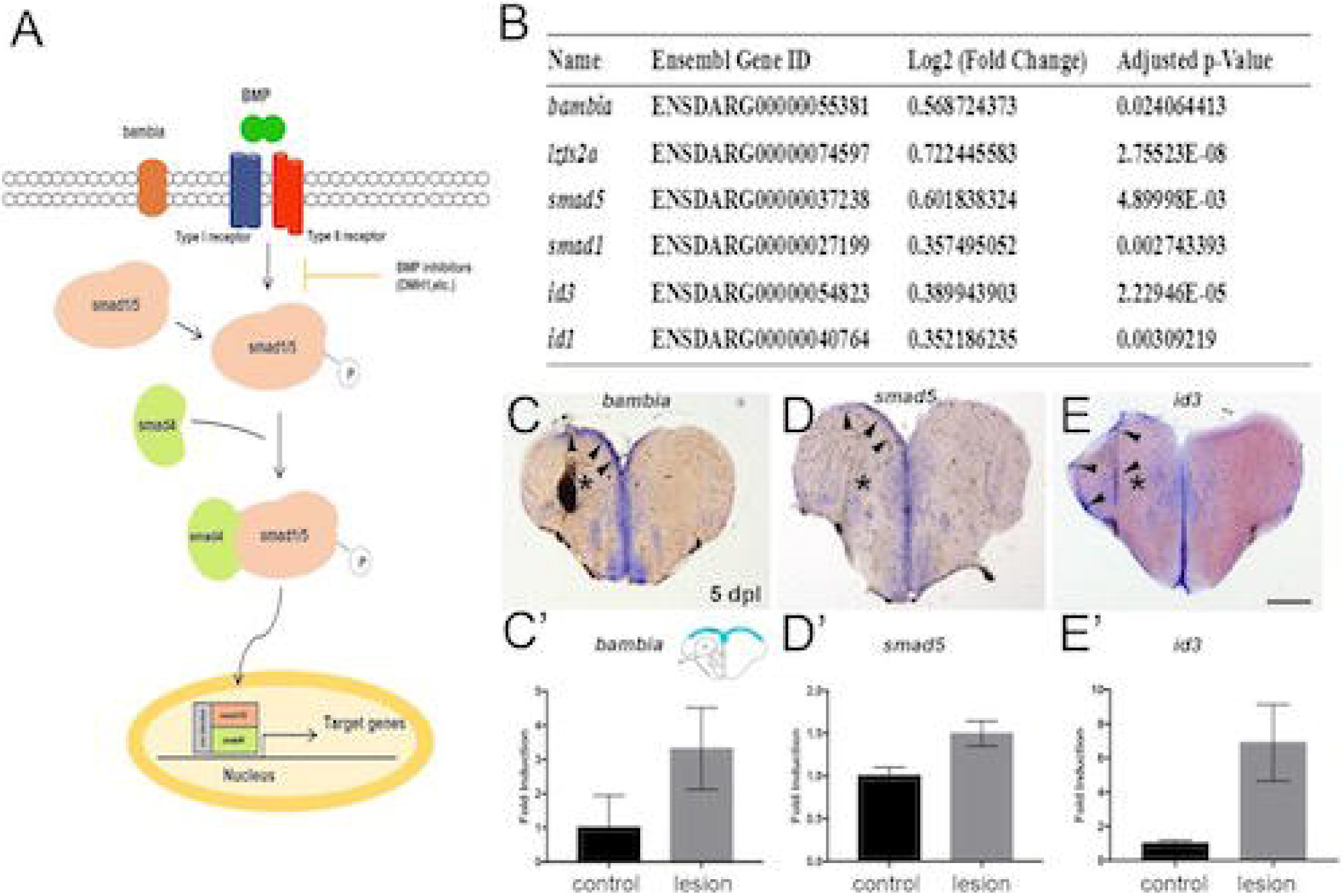
Genes involved in canonical BMP signaling are induced in response to telencephalic injury. (A) Scheme of the BMP signaling pathway. (B) RNAseq analysis of injured telencephala hemispheres in comparison to uninjured hemispheres reveals an up-regulation in mRNA expression of BMP signal transducers (*smad5*), regulators (*bambia*) or downstream target genes (*id3, id1*). (C - E) *In situ* hybridization on sections of the adult zebrafish telencephalon 5 days post lesion. (C) *bambia*, (D) *smad5*, (E) *id3.* Arrowheads indicate up-regulation of gene expression in the left telencephalic hemisphere upon stab injury. (C’-E’) Quantification of (C’) *bambia*, (D’) *smad5* and (E’) *id3* expression up-regulation 5 days post lesion. Only up-regulated areas were quantified along the control and injured ventricular zone from the dorsomedial to the dorsolateral region of the telencephalon (scheme in the upper right-hand corner of (C’) shows the quantified area in green). Scale bar = 200 μm (C, D, E).

**Figure 7:**
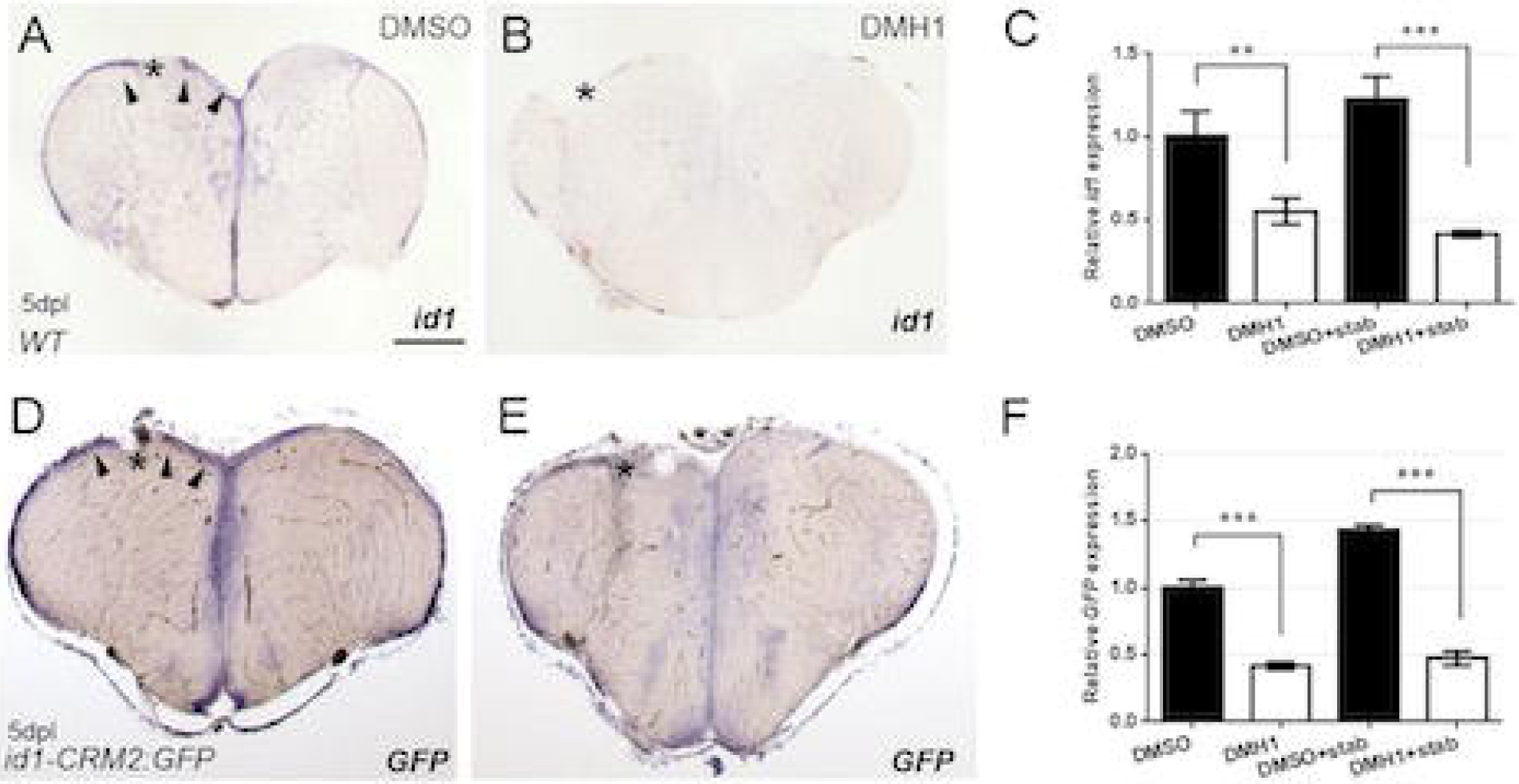
Inhibition of BMP signaling with DMH1 reduces injury induced *id1* and *id1-CRM2:GFP transgene* expression. *ISH* against *id1* (A, B) and *gfp* mRNA (D, E) showed reduction in induction of *id1* (B) and *gfp* (E) transgenesE upon BMP signaling inhibition by DMH1 treatment (B and E) in comparison to DMSO-treated corresponding controls (A and D). Arrowheads indicate up-regulation of gene expression in left telencephalic hemisphere upon stab injury. (C and F) RT-qPCR quantification confirmed reduction in *id1* and *gfp* expression in DMH1 treated fish. The data represent the mean ±SD of three independent experiments. Significance is indicated by asterisks:**, .001≤ *p* <.01; ***, *p* <.001. Scale bar = 200 μm (A, B, D and E). *n* = 3 animals for (C, F).

In summary, our investigation of the transcriptional regulation mediated by *id1-CRM2* suggests that BMP signaling positively regulates the expression of *id1* in the adult zebrafish brain during constitutive and regenerative neurogenesis.

## Discussion

Here, we identified the DNA module *id1-CRM2* as a key regulator of *id1* expression in the ventricular zone of the adult zebrafish telencephalon. Moreover, we show that a *BRE* is crucial for the activity of this CRM and that inhibition of BMP signaling reduces expression of the CRM driven reporter, both during constitutive and reactive neurogenesis. This requirement of BMP signaling is correlated with an increase in mRNAs encoding BMP pathway components and BMP-controlled genes in the transcriptome of injured telencephala.

### BMP as a regulator of *id1* expression during constitutive and regenerative neurogenesis

A key question is which cues control *id1* expression in RGCs during constitutive and regenerative neurogenesis. Inflammatory signals were previously implicated in the induction of regenerative neurogenesis in the zebrafish telencephalon ^43^. However, *id1* expression was not affected by inflammation ^25^. Additionally, Notch signaling, previously shown to be involved in the control of neurogenesis in the zebrafish ^44, 45^, did not affect *id1* expression, either ^25^. Our systematic deletion and mutation analysis of the CRM2 module, as well as pharmacological inhibition of the BMP/Smad signaling pathway strongly suggest that BMP signals are crucial for *id1* expression in the adult zebrafish telencephalon during both constitutive and regenerative neurogenesis. The observation that BMP signaling components (*bmp1aa, smad1, smad5*), as well as known BMP-controlled genes such as *bambia, id3* and *lzts2a* are induced upon brain lesion in addition to the *id1* gene suggests that injury increases the basal level of BMP signaling in the telencephalon.

Remarkably, injury-induced proliferation of NSCs and up-regulation of BMP target genes are always restricted to the injured side of the telencephalon (e.g. ^20, 25^ and this study). Given the close juxtaposition of left and right hemispheres in the medial parts of the telencephalon, this restriction of induction in gene expression and proliferation to the injured half is rather remarkable and implies a highly limited diffusion of the BMP signals towards the uninjured hemisphere. Rather than diffusing freely within the injured side, BMP signals may be sensed and relayed down to the cell bodies at the ventricular zone by the long processes of the RGCs. *Bmp2, bmp4* and *bmp7* mRNAs are expressed in the telencephalon (unpublished data), but the type of BMP signals involved in injury response as well as their origin are unclear. Although, we detected mRNA expression of these *bmps* in our transcriptome data, the change in response to injury was not significant. We did also not observe a reduction in the expression of BMP signaling inhibitors, such as *smad6, smad7, noggin, and follistatin* genes (data not shown). Therefore, it is tempting to speculate that the lesion triggers either release of BMPs or their maturation.

### BMPs as regulators of quiescence and proliferation of neural stem cells in reactive neurogenesis

In agreement with our data, previous work in mouse associated BMP signaling and *id* genes with maintenance of the adult NSCs of the hippocampus and lateral ventricle during constitutive neurogenesis ^46-48^. However, in mice the situation is more complicated as several *id* genes act redundantly ^48^. A single *id* gene appears to provide this function in the zebrafish^25^.

Here, we provide evidence that the mechanism of neural stem cell maintenance by *id1* is conserved not only in constitutive adult neurogenesis, as observed in mouse, but also in the teleost-specific reactive neurogenesis which is involved in injury repair. BMP signals appear to play a crucial role in elevating *id1* expression in response to injury and in this way to eventually down-regulate proliferation of RGCs, securing a pool of resting stem cells for future activation. In this context, it is important to stress that *id1* expression reacts in a delayed fashion relative to the induction of proliferation ^25^. Another signaling system maintaining adult neural stem cell quiescence is the Notch pathway in both zebrafish and mice ^44, 49^. It remains to be assessed whether the BMP/Id1 and Notch/Her4.1 pathways are redundant or parallel pathways with distinct functions in stem cell maintenance during constitutive and reactive neurogenesis. In mouse it was recently suggested that both Notch/Hes and Bmp/Id pathways interact to enhance quiescence of neural stem cells ^50, 51^.

### *Id1-CRM2* is structurally and functionally highly conserved

*Id1-CRM2* is a highly conserved CRM with homologous sequences in all-vertebrate species examined so far (this report, ^52^). The 74 bp central region of *id1-CRM2-core* situated between the *foxA2* and *egr1* binding sites is very similar to a previously identified *BRE* of the mouse and human *id1* regulatory sequence ^33, 34^. The two zebrafish *SBM1* and S*BM2* located in the central region perfectly match the smad binding site consensus GGCGCC (^53, 54^) and that of the *smad binding element (SBE)*, AGAC (^55, 56^). These elements are critical for BMP-induced *id1* expression in the mouse C2C12 myoblast cell line (^33^). Transgenes harboring tandem copies of this central *BRE* served as reliable reporters of canonical BMP signaling activity in mice and zebrafish embryos ^38, 39, 57, 58^.

However, this region was found to be necessary but not sufficient in *id1-CRM2* to drive expression of *id1* in RGCs of the adult zebrafish telencephalon. Moreover, the tandem copy of the BRE did not mediate expression in RGCs of the telencephalon. Conserved *cre* and a *pknox* site were found to be necessary in addition to the *BRE* core of *id1-CRM2* for basal expression in the RGCs. The combination of smad binding sites with *cre* is a conserved feature of the *id1-CRM2*, which is shared with many other BMP target modules in the mammalian genome ^59^. In mouse osteoblasts, a cAMP response element was shown to enhance the response of *id1* to BMP signals ^60^, showing that this interaction is not only restricted to neural stem cells in the zebrafish but is also employed during bone formation in mammals.

The *pknox* binding site in the *id1-CRM2* partially overlaps with the *BRE* sequences (CGCC, CAGC) identified in mouse *id1* to be necessary for strong responsiveness to BMP signaling ^33^. Therefore, it may be possible that these mutations impair the *BRE*. However, the mutations of the *pknox and cre* sites did not impair the BMP mediated response to injury. Our data suggest that their function is dispensable for BMP mediated induction of *id1* reporter expression in response to injury, suggesting that basal and induced expression may involve different co-factors.

Taken together this regulatory region of the *id1* gene appears to be structurally highly conserved between fish and mammals and serves as a regulatory interface that integrates multiple inputs. The high structural conservation is reflected by the fact that the human sequence can drive expression in RGCs of the adult zebrafish telencephalon. Remarkably, this conservation of function is not restricted to constitutive neurogenesis, but the human sequence also faithfully reproduces the response to injury. Thus, despite the vast difference in regenerative capacity, the underlying basic mechanism of reactive neurogenesis appears to be conserved between fish and mammals. This underscores the value of studies in the zebrafish as a model to develop therapies for injuries of the human brain. Clearly, *id1-CRM2* is a highly versatile CRM that drives expression in the zebrafish embryo in various tissues including the notochord (^52^). It is thus expected to contain other elements, some of which may not reveal themselves by conserved sequence homology ^52^.

## Conclusion

Here, we have identified a RGC cis-regulatory module of *id1*, which mediates the input from the BMP signaling pathway into the adult neural stem cells during constitutive and regenerative neurogenesis in the zebrafish telencephalon. This CRM has a high potential to serve as an interface, which will permit to alter the balance between proliferation and maintenance of stem cells in experimental, as well as medical applications.

## Acknowledgments

We thank Nadine Borel and the fish facility staff for fish care, Sabrina Weber for technical support, Maryam Rastegar for her support with the microscopes, and Thomas Dickmeis for discussion and proofreading of the manuscript, Salim Seyfried, Elise Cau, Natascia Tiso and Matthias Hammerschmidt for sharing zebrafish transgenic lines. We are grateful for support by the EU IP ZF-Health (Grant number: FP7-242048), the Deutsche Forschungsgemeinschaft (GRK2039), the programme BioInterfaces in Technology and Medicine of the Helmholtz foundation and the European Union’s Horizon 3952020 research and innovation programme under the Marie Sklodowska-Curie grant agreement No. 643062 (ZENCODE-ITN).

## Role of authors

SR and US: designed the experiments and supervised the work. GZ, MF, LL, ND and TB: conducted the experiments; VG: performed the RNA sequencing data analysis; GZ, MT, ND, SR, US: analyzed the data; SR and US wrote the manuscript.

## Supplementary figure legends

**Supplementary figure 1: The GFP expression pattern of 4 *id1* transgenic lines.** (A) *TgBAC(id1:GFP)*, (B) *Tg(id1-CRM2:GFP)*, (C) *Tg(id1-CRM2-core:GFP)* and (D) Human *Tg(id1-CRM2:GFP*) show a similar expression pattern in the zebrafish embryo at 24 hpf. (A) Annotation of zebrafish embryo structures where GFP is expressed.

**Supplementary figure 2: Relative population sizes of type 1 and type 2 RGCs for the *id1-CRM2* construct at 5dpl, 10dpl and 15dpl**. The Left panel represents relative population sizes of type 1 and type 2 in the unstabbed control telencephalic hemisphere. The right panel represents relative population sizes of type 1 and type 2 in the stabbed injured telencephalic hemisphere. Note that the proportion of GFP^+^ cycling and quiescent RGCs remains constant.

**Supplementary figure 3: The conserved BMP response element in the *id1-CRM2* is crucial for correct expression of the GFP reporter in the RGCs and its induction upon stab wound injury.** Expression of *id1-CRM2-Δ74* (A-D) and *id1-CRM2-mut-SBMs* (E-H) driven reporter constructs is not induced upon stab injury. No GFP^+^/S100^+^ cells can be observed in (D) and (H). Area of magnification is indicated in the upper right-hand corner of each image. Scale bar = 20 μm.

**Supplementary movie 1: Expression of the *id1-CRM2:GFP* transgene in the adult zebrafish ventricular zone.** This expression overlaps perfectly with the expression of the RGC marker S100ß. S100ß expression is shown in blue and *id1-CRM2:GFP* expression in green.

**Supplementary data S1: Sequences of *id1* CRM1/2/3/4/5 compared to orthologus sequences in *Oryzias latipes, Gasterosteus aculeatus* and *Tetraodon nigroviridis* species.** The asterisks at the bottom of the multiple sequence alignments indicate conserved nucleotides in all sequences.

**Supplementary data S2:** Sequences for primers used in this study.

**Graphical abstract legend: The BMP pathway controls *id1* expression in the RGCs both during constitutive and regenerative neurogenesis**.

An evolutionary conserved cis-regulatory module (CRM) of *id1* is necessary for its radial glial cell (RGC) specific expression. The *id1* CRM contains a BMP responsive element (BRE), which is necessary but not sufficient for the expression of *id1* in the ventricular zone of the zebrafish adult telencephalon. The human CRM2 works in zebrafish. Thus its function was conserved during evolution.

