## Supplementary figures and images for "Bone morphogenetic protein signaling regulates Id1 mediated neural stem cell quiescence in the adult zebrafish brain via a phylogenetically conserved enhancer module"

### Supplemental Figure3

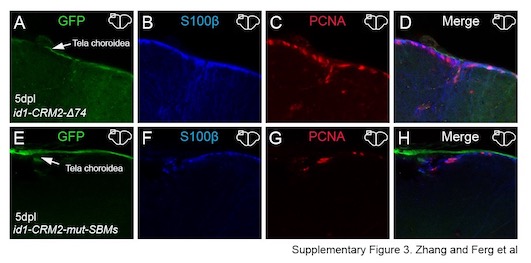

### Supplemental Figure 1

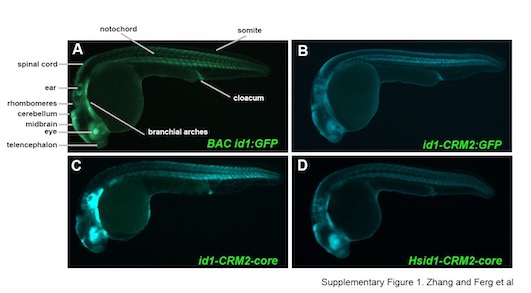

### Supplemental Figure 2

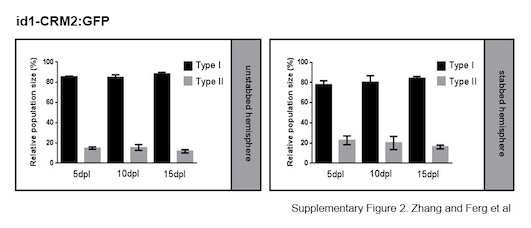
